# Strategies for Inducing and Validating Zinc Deficiency and Zinc Repletion

**DOI:** 10.1101/2024.02.28.582542

**Authors:** Tara-Yesomi Wenegieme, Dalia Elased, Kelia E. McMichael, Jananie Rockwood, Khanzada Hasrat, Adaku C. Ume, Andrea G. Marshall, Kit Neikirk, Annet Kirabo, Khalid M. Elased, Antentor Hinton, Clintoria R. Williams

## Abstract

Given the growing interest in the role of zinc in the onset and progression of diseases, there is a crucial demand for reliable methods to modulate zinc homeostasis. Using a dietary approach, we provide validated strategies to alter whole-body zinc in mice, applicable across species. For confirmation of zinc status, animal growth rates as well as plasma and urine zinc levels were evaluated. The accessible and cost-effective methodology outlined will increase scientific rigor, ensuring reproducibility in studies exploring the impact of zinc deficiency and repletion on the onset and progression of diseases.

**New and Noteworthy:** This methods paper details 1) dietary approaches to alter zinc homeostasis in rodents, and 2) qualitative and quantitative methods to ensure the zinc status of experimental animals. The outlined accessible and cost-effective protocol will elevate scientific rigor, ensuring reproducibility in studies exploring the impact of zinc deficiency and repletion on the onset and progression of a multitude of health conditions and diseases.

**Graphical Abstract:** 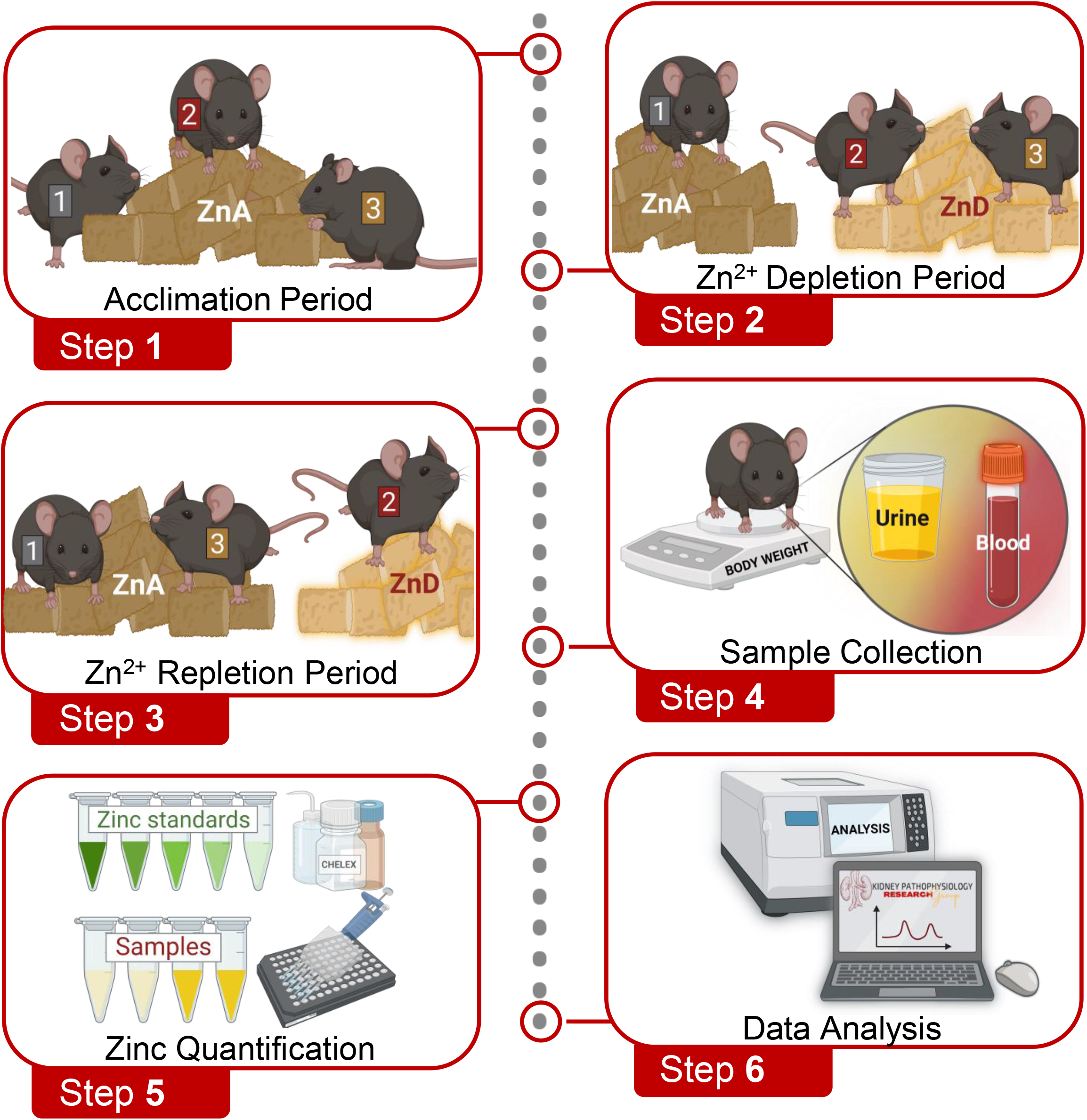

## Introduction

Zn^2+^ is the second most abundant trace metal in the human body, constituting 2 to 3 grams^1^. Skeletal muscle and bone are major Zn^2+^ reservoirs, accounting for approximately 50% and 30% of total body Zn^2+^, respectively^2–5^. Lower Zn^2+^ fractions are distributed across various tissues, including the kidney, prostate, liver, gastrointestinal tract, skin, lung, brain, heart, and pancreas^5–9^. Within these organs, intracellular Zn^2+^ serves as an essential cofactor for the catalytic activity and structural integrity of over 300 proteins, including those involved in synthesis of macromolecules and cell division^10–13^.

Since its initial recognition in 1963^14^, Zn^2+^ deficiency has been increasingly implicated as a hidden culprit in multiple health conditions and chronic diseases such as diabetes, kidney diseases, cancers, neurodegenerative diseases, gastrointestinal disorders, respiratory infections, and skin conditions^5,15– 17^. For more than 60 years, epidemiological, clinical, and experimental studies have focused on determining the role of Zn^2+^ deficiency in the development of these conditions and diseases^14,17–20^. Given the continued interest in the role of zinc in the onset and progression of diseases, more research is needed to identify the impact of this important micronutrient. The knowledge gained from these valuable studies will advocate for effective approaches that integrate Zn^2+^ supplementation into existing therapeutic strategies that address a plethora of conditions and diseases.

To directly study the impact of this important micronutrient on health and disease, standardized methods are required to modulate zinc homeostasis. Using a dietary approach, we provide validated strategies to alter whole-body zinc in mice, applicable across species. We also provide qualitative and quantitative methods to ensure the zinc status of experimental animals. The outlined accessible and cost-effective protocol will elevate scientific rigor, ensuring reproducibility in studies exploring the impact of zinc deficiency and repletion on the onset and progression of a multitude of health conditions and diseases.

## Table of Key Resources

| REAGENT or RESOURCE | SOURCE | IDENTIFIER |
| --- | --- | --- |
| **Chemicals** |  |  |
| Chelex 100 Chelating Resin | Bio-Rad, Hercules, CA | 1421253 |
| Tris Buffer Saline Table | Millipore-Sigma, MO, USA | T5030-100TAB |
| Protease Inhibitor Cocktail | Millipore-Sigma, MO, USA | P8340-1ML |
| **Critical Commercial Assays** |  |  |
| Zinc Quantification Kit | Abcam, Boston, MA | ab176725 |
| **Experimental Models** |  |  |
| C57BL/6J Male Mice | The Jackson Laboratory | JAX:000664 |
| **Other** |  |  |
| Standard Chow Diet | Teklad, Inotiv, IN, USA | Teklad 2018 |
| Zn<sup>2+</sup> Adequate Diet (50 ppm) | Teklad, Inotiv, IN, USA | TD 85420 |
| Zn<sup>2+</sup> Deficient Diet (1 ppm) | Teklad, Inotiv, IN, USA | TD 85419 |
| Deionized Distilled H<sub>2</sub>O | Millipore-Sigma, MO, USA | 6442-88 |
| Purified Drinking H<sub>2</sub>O | Aquafina, Sam's Club | 988708 |
| Black, Clear Bottom 96-well Plate | Thermo Fisher Scientific, Rochester, NY | 265301 |
| 15 ml Conical Tubes (Clear) | Celltreat, Pepperell, MA | 230411-R |
| 15ml Conical Tubes (Black) | Celltreat, Pepperell, MA | 229431 |
| 50 ml conical tube (Clear) | Thermo Fisher Scientific, Rochester, NY | 14-432-22 |
| Microcentrifuge Tubes (Black) | Celltreat, Pepperell, MA | 229437 |
| Microcentrifuge Tubes (Clear) | Electron Microscopy Sciences, Hatfield, PA | 63027-02 |

## Materials and Equipment

### 4% Chelex Solution

- Suspend Chelex 100 Chelating Resin (2 g) in double distilled H_2_0 (50 mL).
- On a rocker, shake Chelex Solution for 2 - 3 minutes.
- Centrifuge Chelex Solution at 6,000rpm for 1.5 - 2 minutes.
- Collect clear portion of supernatant and aliquot 10 mL into multiple clear conical tubes.
- Store 4% Chelex Solution aliquots at -80° C until use.

### Tris Buffer

- In an aliquot of 4% Chelex Solution (10 mL), suspend a Tris Buffered Saline Tablet.
- Using a rocker, gently agitate the buffer until the tablet is dissolved.
- Aliquot Tris Buffer (5 mL) into black conical tubes.
- Store Tris Buffer aliquots at 4°C until use.

### Zn Detector Solution

- Into an aliquot of Tris Buffer (5 mL), dispense Zn Detector (25 μl). **Note**: The Zn Detector is light sensitive.
- Vortex for 10 seconds.
- Immediately use Zn Detector Solution for zinc quantification.

### ZnCl_2_ Standards (Table 1)

- Using reagents provided in the Zinc Quantification Kit, prepare a 1 mM ZnCl_2_ Standard by suspending 100 mM ZnCl_2_ Standard (10 μL) into Assay Buffer (990 μL).
- To prepare a 100 μM ZnCl_2_ Standard, suspend 1 mM ZnCl_2_ (100 μL) into Tris Buffer (900 μL).
- To prepare the designed concentrations of ZnCl_2_ Standards, begin with the 100 μM ZnCl_2_ Standard and perform a serial dilution in Tris Buffer.
- Store ZnCl_2_ Standards at 4°C until use. **Note**: Discard ZnCl_2_ Standards after two uses.

**Table 1.**

| **Table 1** |  |  |  |  |
| --- | --- | --- | --- | --- |
| **ZnCl<sub>2</sub> Standards** |  | **Tris Buffer** |  | **ZnCl<sub>2</sub> Concentration** |
| 100 µL of 1 mM | + | 900 µL | = | 100 µM |
| 500 µL of 100 µM | + | 500 µL | = | 50 µM |
| 500 µL of 50 µM | + | 500 µL | = | 25 µM |
| 500 µL of 25 µM | + | 500 µL | = | 12.5 µM |
| 500 µL of 12.5 µM | + | 500 µL | = | 6.25 µM |
| 500 µL of 6.25 µM | + | 500 µL | = | 3.125 µM |
| 500 µL of 3.125 µM | + | 500 µL | = | 1.56 µM |
| 500 µL of 1.56 µM | + | 500 µL | = | 0.78 µM |
| 500 µL of 0.78 µM | + | 500 µL | = | 0.39 µM |
| 500 µL of 0.39 µM | + | 500 µL | = | 0.195 µM |
| 500 µL of 0.195 µM | + | 500 µL | = | 0.097 µM |
| ----- | + | 1000 µL | = | 0 µM |

## Before you Begin

### Institutional permissions

All animal experiments performed were in accordance with the Animal Care and Use Committee of Wright State University and were conducted in accordance with the National Institutes of Health Guide for the Care and Use of Laboratory Animals. The appropriate licenses and permission for animal experiments should be obtained before proceeding with this protocol.

## Step-by-Step Method Details

I. **Acclimation Period** **Timing: [2 weeks]** **<u>Acclimation of Mice to Zn</u>**^**<u>2+</u>**^**<u>-Adequate Diet</u>**
  - **CRITICAL**: Group house mice at 27°C in a 12:12-hour light-dark cycled room.
    a. Until 6 weeks of age, maintain C57BL/6 mice on a standard chow diet.
    b. At 6 weeks old, acclimate mice to a Zn^2+^-adequate (ZnA) diet for 2 weeks.
      i. Provide mice with ad libitum access to food and purified drinking H_2_O. **NOTE**: Since the quality of tap H_2_O may vary between facilities, use commercially available purified drinking H_2_O.
    c. Each week, weigh individual mice to monitor growth rate.
    d. At the end of the acclimation period, proceed to “Step-by-Step Method Details” section.
II. **Zn**^**2+**^ **Depletion Period** **Timing: 10 weeks**
  - **CRITICAL:** Since mice are social animals, group housing provides a less stressful environment than individually housing mice. If mice must be housed individually, ensure that all experimental groups are housed similarly. To ensure that mice consume similar nutritional content, pair-feed mice by providing the ZnA group with the amount consumed by ZnD mice. Avoid using glass H_2_O bottles, as glass may be a source of Zn^2+^. To avoid consumption of fecal Zn^2+^, weekly mouse cage changes are required.
    a. **<u>Group 1 (Zn</u>**^**<u>2+</u>**^ **<u>Adequate):</u>** For the control group, maintain a subset of the ZnA mice on the ZnA diet for up to 10 weeks. Continue to provide ZnA mice with commercially available purified drinking H_2_O.
    b. **<u>Group 2 (Zn</u>**^**<u>2+</u>**^ **<u>Deficient):</u>** Switch a subset of ZnA mice to an ingredient-matched, ZnD diet for 10 weeks. To maintain a Zn^2+^-free environment, provide ZnD mice deionized, distilled drinking H_2_O in plastic containers. **CRITICAL:** Visual signs of chronic Zn^2+^ deficiency include growth retardation (**Figure 1B**), hair loss (**Figure 1C**) and loose stool. However, the absence of these visual signs should not disqualify mice from studies. Zn^2+^ levels must be quantified to verify changes in Zn^2+^ levels (see “Zn^2+^ Quantification” Step).

**Figure 1:**
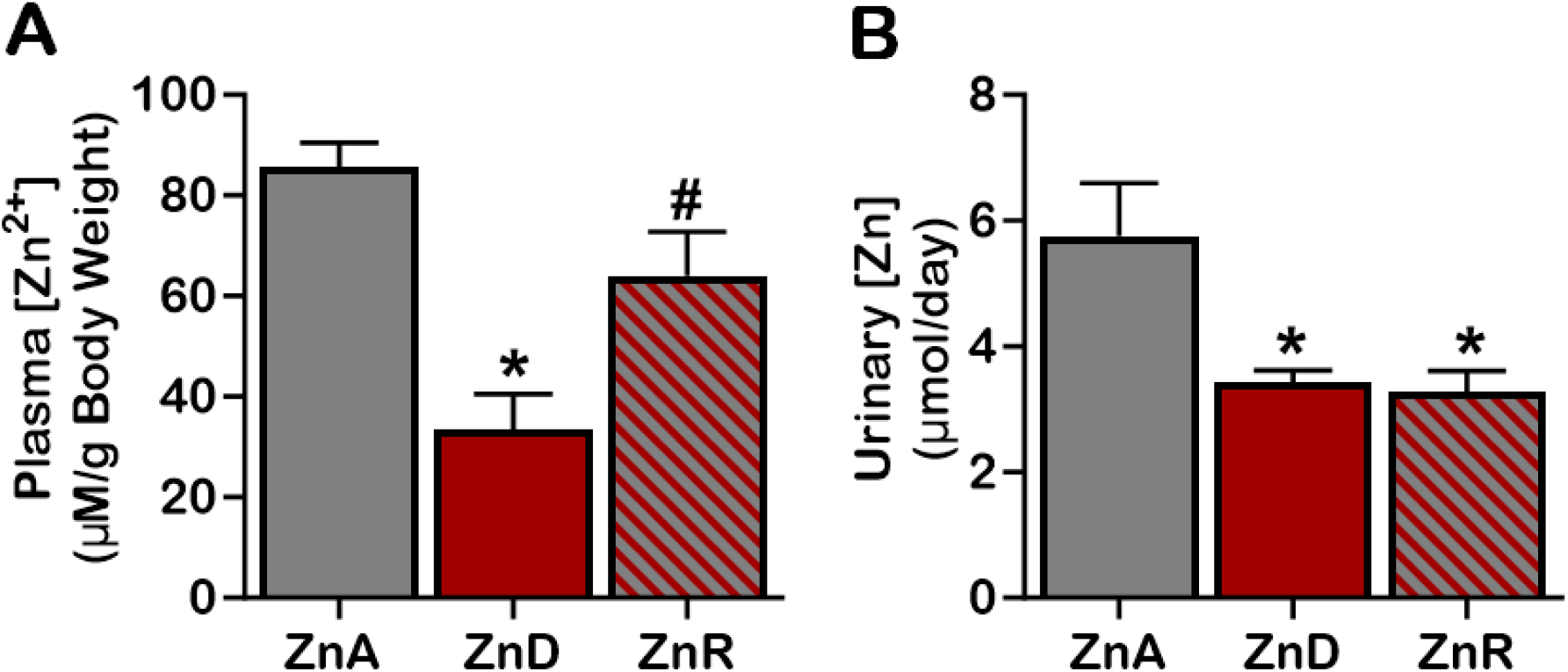
Zn^2+^ homeostasis plays a critical role in growth and development. To investigate the impact of dietary Zn restriction on mouse health, mice were fed a Zn^2+^-adequate diet (ZnA, grey) or Zn^2+^-deficient diet (ZnD, red) for 10 weeks. Effects on weekly weight gain (**Figure 1A)**, body mass at week-10 (**Figure 1B**), and skin health (**Figure 1C**) were examined. *p < 0.05 vs ZnA, ^#^p < 0.05 vs ZnD. **NOTE:** While Zn^2+^ deficiency was confirmed at week-10 (**Figure 2**), this status may occur at an earlier timepoint. As such, plasma and urine Zn^2+^ levels should be quantified to identify timepoint of changes in Zn^2+^ levels (see “Zn^2+^ Quantification” Step).

**Figure 2:**
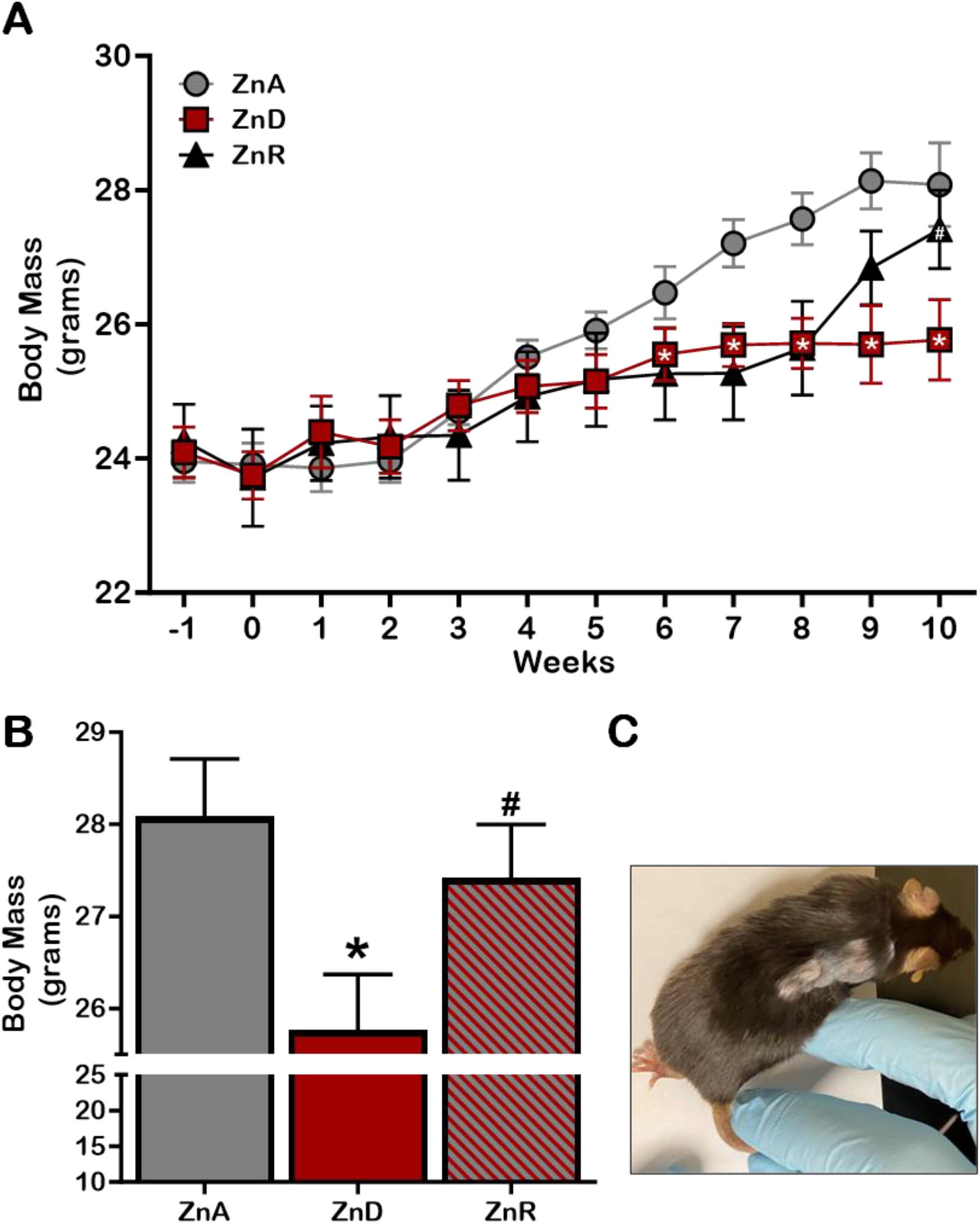
Impact of dietary Zn^2+^ restriction and repletion on circulating Zn^2+^ levels. To probe the influence of dietary Zn restriction on Zn^2+^ levels, mice were fed a Zn^2+^-adequate diet (ZnA, grey) or Zn^2+^-deficient diet (ZnD, red) for 10 weeks. Effects on plasma (**Figure 1A**), and urine (**Figure 1A**) Zn^2+^ levels were assessed. *p < 0.05 vs ZnA, ^#^p < 0.05 vs ZnD.
III. **Zn**^**2+**^ **Repletion Period** **Timing: 2 weeks**
  c. **<u>Group 3 (Zn</u>**^**<u>2+</u>**^ **<u>Repletion):</u>** For Zn^2+^ repletion studies, after 8 weeks on a ZnD-diet (as described for Group 2), return mice to the ZnA-diet and purified drinking H_2_O (as described for Group 1). For the remaining 2 weeks of the study, maintain this subset of mice (ZnR) on the ZnA-protocol.
IV. **Sample Collection**
  a. **Plasma Samples**
    1. From the submandibular vein, collect blood samples from mice. **NOTE**: Collection method should be perfected to avoid hemolysis.
    2. Centrifuge collected blood samples at 1500 x g at 4 °C for 15 minutes.
    3. Collect and aliquot plasma (20 μL) into clear microcentrifuge tubes.
    4. Proceed to “Zn^2+^ Quantification” Step.
      - Optional Step: Store plasma samples at -80°C for up to 12 months.
  b. **Urine Samples**
    1. To collect 24-hour urine samples, house mice in metabolic cages.
    2. Centrifuge collected urine samples at 2000 x g at 4 °C for 10 minutes.
    3. Collect and aliquot debris-free urine (100 μL) into black microcentrifuge tubes.
    4. Proceed to “Zn^2+^ Quantification” Step.
      - Optional Step: Store urine samples at -80°C for up to 12 months.
V. **Zinc Quantification** **Timing: [30 mins]** **NOTE**: Perform all steps in a dark area.
  a. Dispense Tris Buffer (30 μl) into appropriate wells of a black, clear bottom 96-well plate.
  b. In duplicate, pipet Zn^2+^ Standards (10 μl) and unknown samples (10 μl) into the appropriate wells of the 96-well plate.
  c. To protect Zn^2+^ Standards and unknown samples from light, cover 96-well plate with aluminum foil.
  d. Using a rocker, gently shake 96-well plate at room temperature for 15 minutes to ensure mixing of Zn^2+^ Standards and unknown samples with Tris Buffer.
  e. With a fluorescence plate reader, measure background fluorescence (Ex/Em = 485/525) of Zn^2+^ Standards and unknown samples.
  f. Dispense Zn Detector (30 μl) into the appropriate wells of the 96-well plate.
  g. After covering the 96-well plate with aluminum foil, gently shake on a rocker for 10 seconds.
  h. Incubate covered 96-well plate at room temperature for 15 minutes and then measure Zn Detector fluorescence (Ex/Em = 485/525).
VI. **Data Analysis**
  a. To obtain net fluorescence, subtract background fluorescence from Zn Detector fluorescence.
  b. Using the net fluorescence, calculate Zn^2+^ concentration of unknown samples by extrapolating values from the Zn^2+^ standard curve.

## Expected Outcomes

### Impact of Dietary Zn^2+^ Restriction and Repletion on Growth and Development

To assess the impact of Zn^2+^ restriction on nutritional status, body mass was measured in mice placed on a ZnA or ZnD-diet for 10 weeks (**Figure 1**). Until week-5, mice on a ZnD diet had similar body masses as mice maintained on the ZnA diet (**Figure 1A**). However, starting at week 6, ZnD mice experienced slower weight gain. Furthermore, between 6 - 8 weeks, some ZnD mice experienced alopecia (**Figure 1C**). While average food consumption was not significantly different (14.74 g ± 0.21 vs 15.21 g ± 0.54), at week-10, mice on the ZnD diet had significantly less body mass (25.769 g ± 0.601) compared to ZnA mice (28.08 g ± 0.626) (**Figure 1B**).

To assess the impact of Zn^2+^ repletion (ZnR) on nutritional status, at week 8, ZnD mice were returned to a ZnA diet. Zn^2+^ repleted mice experienced an accelerated growth rate (**Figure 1A**). After 2 weeks, body mass (27.42 g ± 0.583) and food consumption (15.78 g ± 0.39) of ZnR mice (**Figure 1B**) was comparable to mice maintained on the ZnA diet. *These results indicate that chronic dietary Zn*^*2+*^ *restriction impairs growth, and acute dietary Zn*^*2+*^ *repletion improves development*.

### Impact of Dietary Zn^2+^ Restriction and Repletion on Plasma and Urinary Zn^2+^ Levels

Using the Zn^2+^ Quantification protocol, plasma Zn^2+^ levels were quantified and normalized to mouse body mass (**Figure 2A**). At week-10, mice maintained on the ZnA-diet had plasma Zn^2+^ levels of 3.19 μM/g Body Mass. In contrast, plasma Zn^2+^ levels were 40% lower in mice placed on a ZnD diet (1.29 μM/g Body Mass). For ZnD mice returned to the ZnA-diet (ZnR) for 2 weeks, plasma Zn^2+^ levels (2.44 μM/g Body Mass) increased and were comparable to mice maintained on the ZnA diet.

The impact of dietary Zn^2+^ restriction and repletion on urinary Zn^2+^ excretion was also examined (**Figure 2B**). Similar to plasma Zn^2+^, urinary Zn^2+^ levels were reduced in ZnD mice (3.43 μmol/day ± 0.19 vs 5.76 μmol/day ± 0.84). However, after 2 weeks of Zn^2+^ repletion, urinary Zn^2+^ levels remained low (3.27μmol/day ± 0.34) and were comparable to mice maintained on the ZnD diet. *Collectively, these findings demonstrate that chronic dietary Zn*^*2+*^ *restriction is effective at inducing Zn*^*2+*^ *deficiency. Furthermore, while acute dietary Zn*^*2+*^ *repletion improves circulating Zn*^*2+*^ *levels, urinary Zn*^*2+*^*excretion is not yet restored*.

## Limitations

This protocol was developed and optimized in adult, male, wild-type mice on a C57Bl/6 background. Various factors influence the handling of Zn^2+^, including genetic composition, age, and sex. Given the potential diversity in how Zn^2+^ is handled, the impact of dietary Zn^2+^ depletion and repletion may vary between experimental animals within a study. Consequently, the intricate interplay of genetic factors, age-related dynamics, and sex-specific differences can contribute to distinct responses to Zn^2+^ manipulation. This important consideration underscores the significance of establishing baseline Zn^2+^ levels. By obtaining a reference point, Zn^2+^ changes can be assessed not only between different experimental groups but also longitudinally, with each animal also serving as its own control. This rigorous approach provides a comprehensive assessment of the impact of Zn^2+^ dynamics on physiological and pathophysiological processes.

## Troubleshooting

### Problem 1: No changes in plasma and/or urinary Zn^2+^ levels after Zn^2+^ depletion

#### Potential solution: Extend duration of Zn^2+^ Depletion Period

- To maintain whole-body Zn^2+^ homeostasis, Zn^2+^ is released from Zn^2+^ stores into the circulation. As such, a longer depletion period may be required to deplete Zn^2+^ stores and subsequently reduce serum and urine Zn^2+^ levels.

### Problem 2: Reduced plasma and urinary Zn^2+^ levels after Zn^2+^ repletion

#### Potential Solution: Extend duration of Zn^2+^ Repletion Period

- While ZnR mice (2.44 μM/g Body Mass) have increased plasma Zn^2+^ levels compared to ZnD mice (1.29 μM/g Body Mass), these levels are not yet comparable to ZnA mice (3.19 μM/g Body Mass). Furthermore, urinary Zn^2+^ levels are still depleted (3.27 μmol/day ± 0.34 vs 5.76 μmol/day ± 0.84). Repletion of Zn^2+^ stores within organs proceed repletion of serum and urinary Zn^2+^ levels. As such, a longer repletion period may be required when using serum and urine Zn^2+^ as markers of Zn^2+^ status.

## Resource availability

### Lead contact

*Further information and requests for resources and reagents should be directed to and will be fulfilled by the lead contact, Clintoria R. Williams, PhD, FAHA (clintoria*.*williams@wright*.*edu)*.

### Materials availability

*There are no newly generated materials associated with this protocol*.

### Data and code availability

*This study did not generate/analyze [datasets/code]*.

## Acknowledgements

This work was supported by the National Institute of Diabetes and Digestive and Kidney Diseases Grants: R21 DK119879 (to C.R.W.), and R01 DK-133698 (to C.R.W.); by the American Heart Association Grant 16SDG27080009 (to C. R.W.); and by the American Society of Nephrology KidneyCure Transition to Independence Grant (to C.R.W.).

## Author contributions

T.Y.W., K.E.M., D.E.K., C.R.W., performed the experiments; T.Y.W., K.E.M., J.R., and C.R.W. analyzed the data; T.Y.W., K.E.M., J.R., and C.R.W. interpreted the results of the experiments; T.Y.W., K.E.M., J.R., A.C.U., and C.R.W. prepared the figures; T.Y.W., K.E.M., J.R., K.H., A.M., A.H., A.K., K.N., and C.R.W. drafted the manuscript; T.Y.W., J.R. and C.R.W. edited and revised the manuscript; T.Y.W, J.R., and C.R.W. approved the final version of the manuscript.

